# smalldisco, a pipeline for siRNA discovery and 3’ tail identification

**DOI:** 10.1101/2022.07.15.500275

**Authors:** Ian V. Caldas, Leanne H. Kelley, Yasir H. Ahmed-Braimah, Eleanor M. Maine

## Abstract

Capturing and sequencing small RNAs is standard practice, however identification of a group of these small RNAs—small interfering RNAs (siRNAs)—has been more difficult. We present smalldisco, a command-line tool for *<u>small</u>* interfering RNA *<u>disco</u>*very and annotation from small RNA-seq datasets. smalldisco can distinguish short reads that map antisense to an annotated genomic feature (e.g., exons or mRNAs), annotate these siRNAs, and quantify their abundance. smalldisco also uses the program Tailor to quantify 3’ non-templated nucleotides of siRNAs or any small RNA species. smalldisco and supporting documentation are available for download from GitHub (https://github.com/ianvcaldas/smalldisco).

## Introduction

Small, non-coding RNAs (sRNAs) are critical components of transcriptional and post-transcriptional gene regulatory mechanisms in plants, animals, and fungi (Castel and Martienssen 2013, Holoch and Moazed 2015, Ozata *et al*. 2019, Galagali and Kim 2020, Ketting and Cochella 2021). Three abundant types of sRNAs — micro (mi) RNAs, PIWI-associated (pi) RNAs, and small interfering (si) RNAs — associate with Argonaute proteins and provide specificity via complementary base pairing with target RNAs (Iwakawa and Tomari 2022). Although the details of sRNA biogenesis vary among species, distinct miRNA and piRNA gene clusters are present in the genome, often under control of dedicated promoters, and expression of these genes produces populations of identical miRNA or piRNA sequences. In contrast, siRNAs in most organisms are generated by Dicer cleavage of double strand (ds) RNA, itself often generated by RNA-directed RNA polymerase (RdRP) activity on single strand (ss) RNA templates. One notable exception is nematodes, where most siRNAs are produced as short RdRP products that are not processed by Dicer. Complicating matters, biogenesis of different sRNA types can be linked, e.g., plants produce phased siRNAs downstream of miRNA-AGO activity on RNA pol II transcripts, and nematodes produce siRNAs downstream of piRNAs. Because miRNAs and piRNAs are identified as discrete, genome-encoded sequences, it is relatively straightforward to identify miRNA and piRNA sequences in sRNA-seq data sets by mapping reads to their canonical loci in the genome. In contrast, siRNA sequences often map to numerous positions across the template RNA and are not always straightforward to identify.

The main methods used for identifying siRNAs have included: (i) recovering sRNAs that co-immunoprecipitate with a specific Argonaute protein (e.g., Claycomb *et al*. 2009, Chaves *et al*. 2021, de Albuquerque *et al*. 2015; Mi *et al*. 2008; Seroussi *et al*., 2022; Svendsen *et al*. 2019, Xu *et al*., 2018; Okamura *et al*., 2013; Gartland *et al*., 2022; Pisacane and Halic 2017); (ii) a differential expression approach to define populations of siRNAs by comparing sRNAs in wildtype and certain mutant backgrounds (e.g., Gu *et al* 2009; Conine *et al*. 2010; Montgomery *et al*. 2008; Hua *et al*. 2021; Krzyszton and Kufel 2022); and (iii) computationally predicting siRNAs from sRNA sequence data (e.g., Montgomery *et al*. 2012, Wei *et al*. 2019, Rogers and Phillips 2020, Scheer *et al*. 2021). Differential expression and AGO immunoprecipitation have been very useful for parsing biogenesis mechanisms and linking individual Argonaute proteins to regulation of certain target genes via specific siRNA guides, although they do not provide a comprehensive view of all siRNA sequences in a transcriptome or their abundance. Computational pipelines have been very useful, but many of those developed for published studies are not being actively maintained nor were developed for general use across different datasets. We seek to overcome this limitation by developing and actively maintaining a user-friendly pipeline for identifying siRNAs within full sRNA-seq data sets.

SiRNA biogenesis varies among organisms, and we chose to identify siRNAs based on pattern rather than biogenesis mode. In other words, although most siRNAs are produced from dsRNA precursors in most organisms, the strand that is antisense to the target RNA is what guides the Argonaute protein to that target. Consequently, we developed smalldisco, a *<u>small</u>*-interfering RNA *<u>disco</u>*very program, for parsing sRNA-seq data sets to identify siRNAs that are antisense to genomic features of interest to the user, such as protein-coding regions. While this approach is most obviously appropriate for *C. elegans*—where most siRNAs are direct RdRP products—we argue that it can provide useful information for other species, as we demonstrate below. Our goal was to develop a broadly applicable pipeline that is useful for addressing various biological questions. After identification, smalldisco can feed its discovered siRNAs into the Tailor program (Chou *et al*. 2015) to identify 3’ non-templated nucleotide additions to these siRNA sequences and produce output in a user-friendly format as tables of siRNA targets, their tail modifications, and their abundances. While smalldisco does not identify miRNAs or piRNAs (programs available for those purposes include unitas [Gebert *et al*. 2017], miRDeep* [An *et al*. 2013], and piPipes [Han *et al*. 2015]), it does provide a wrapper option to identify 3’ tailing modifications to miRNA and piRNA sequences from the same sRNA dataset through Tailor.

## Methods

smalldisco users will need the following files: a folder of BAM files for their samples of sRNA-seq data; a reference genome (FASTA); and an annotated (GTF or GFF) file of the genome of interest. Note that the GTF/GFF file must in be the standard tab-separated, nine-column format (https://useast.ensembl.org/info/website/upload/gff.html). A summary of the pipeline is diagrammed in Figure 1. smalldisco can be run in two modes: `sirna` and `tail`.

**Figure 1.**
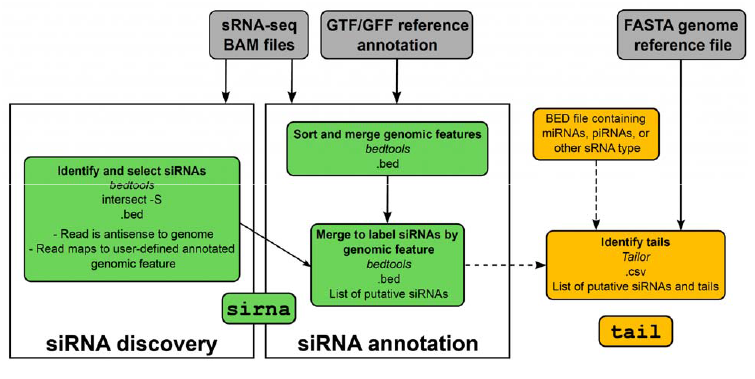
Overview of the smalldisco pipeline outlining the siRNA discovery and annotation workflows. Two analysis modes are available: `sirna` and `tail`. “Genomic feature” refers to a user-defined annotated genomic feature. Dashed arrows indicate only one of these inputs is necessary.

The `sirna` mode implements smalldisco’s siRNA discovery. First, the program creates a set of candidate genomic regions that could contain the siRNAs of interest. These are created from a specified annotated genomic feature, such as exons or 3’ UTRs; overlapping features on the same strand are merged, while any regions of overlap across different strands are removed from the analysis. Second, reads are filtered to those mapping antisense to one of these candidate regions. Third, groups of overlapping reads are merged into a putative siRNA region if they fulfill two criteria: if there are more than r overlapping reads and if the putative siRNA would span s or more base pairs. Both parameters are set by the user, with default r=10 and s=10. The result is a BED file of putative siRNA regions. The handling of siRNA sequences spanning a splice junction in an exon-exon boundary depends on the alignment of reads. If there are antisense reads mapping to both exons, two putative siRNA regions will be reported, while if the reads map to only one of the exons, only one region in that exon will be reported. In the example datasets that we list below (Table S1), 0.01-1.5% of reads within a BAM file are sequences that span an exon-exon junction. Therefore, we recommend that users perform their read alignment with a splice-aware aligner such as HISAT2.

Another consideration is the detection of siRNA regions within transposable elements. If the user chooses to find siRNAs within exons, those would include transposase genes present in intact transposable elements. Repetitive sequences within intact or partial transposable elements could also be defined as the features producing candidate regions (but see the Readme for considerations). Given the complications associated with mapping repetitive element-derived reads to a genome, we recommend that the user map the sRNA reads to a FASTA file of consensus sequences of transposable elements that are derived from their genome of interest.

`tail` mode, the second and optional step of the pipeline, can quantify read tails for the sRNAs. A BED file of sRNA regions, which can be the list of putative siRNA regions generated by the previous step or any other predefined list, is used as input to Tailor (https://github.com/jhhung/Tailor). Tailor maps the reads to the reference genome and adds a tag at the end of each read’s alignment record to indicate the length and sequence of a tail, or to indicate the absence of a tail, producing a tagged BAM file. Tailor implements a default of zero mismatches in the templated sRNA sequence before searching for non-templated tails. Tagged reads are then mapped to the provided sRNA regions file and the tail tags are converted to nucleotides. The output is a CSV file of sRNA targets and tails. In addition to “untailed” and the tail types in the table, one row per target will include a total for “all_tails.” The sum of “untailed” and “all_tails” equals the total number of reads for that target.

We validated smalldisco using publicly available datasets published by Xu *et al*. (2018) (PRJNA447866), Claycomb *et al*. (2009) (PRJNA123485), Scheer *et al*. (2021) (PRJNA624174), and Davis *et al*. (2018) (PRJNA398116). Samples used for each analysis are listed in Table S1. To analyze these datasets, FASTQ files were obtained from NCBI’s Gene Expression Omnibus, trimmed of 3’ adapters using cutadapt, mapped to a reference genome using HISAT2, and run through smalldisco (both modes) with default parameters. For Xu *et al*. and Claycomb *et al*. datasets, the smalldisco siRNA output was normalized to counts per million (CPM) and filtered to retain only siRNA-associated genes with ≥ 25 reads per million, the threshold used by these authors. To normalize the siRNA library as executed by Scheer *et al*., we created a library by counting reads that map to exons, combined this library with the smalldisco siRNA output, and normalized to CPM. Finally, to benchmark smalldisco’s performance against another sRNA discovery tool, segmentSeq (Hardcastle *et al*. 2011), we analyzed two samples from Davis *et al*. (2018), each with three replicates (SRR5931601, SRR5931602, SRR5931603, SRR5931604, SRR5931605,

SRR5931606), and compared runtime, number of identified siRNA “loci,” and resolution of siRNA region sizes.

## Results and discussion

We validated the siRNA discovery function of smalldisco using publicly available datasets from two studies where *C. elegans* Argonaute-specific small RNAs were recovered by immunoprecipitation. Xu *et al*. (2018) immunoprecipitated WAGO-4 Argonaute and recovered siRNAs (present at or above 25 CPM) that corresponded to 4756 target genes. smalldisco analysis of their immunoprecipitated sRNA dataset identified siRNAs corresponding to 4706 target genes using a 25 CPM cut off, including ~93% of those identified by Xu *et al*. (4413) (Figure 2A). Claycomb *et al*. (2009) immunoprecipitated CSR-1 Argonaute and identified siRNAs corresponding to 4147 target genes present at or above 25 CPM and enriched ≥ two-fold in the IP compared to input RNA. smalldisco identified 4209 siRNA target genes in this dataset, including ~81% of those reported by Claycomb *et al*. (2009) (Figure 2B). For both datasets, some sequences were identified as siRNAs only by smalldisco or only by the authors, and we speculate these reflect differences in filtering criteria or data processing. Overall, our results demonstrate that smalldisco can accurately identify siRNAs with known Argonaute association.

**Figure 2.**
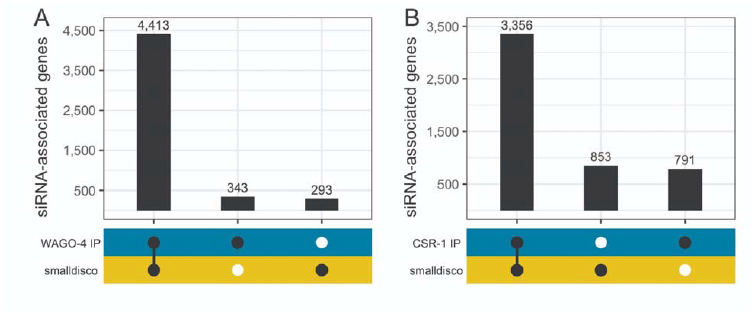
siRNA identification among sRNAs that co-immunoprecipitate with Argonaute proteins. Small RNAs enriched in (A) WAGO-4 immunoprecipitate (Xu *et al*. 2018) and (B) CSR-1 immunoprecipitate (Claycomb *et al*. 2009) were analyzed with smalldisco and the output was compared to the authors’ reported findings. Plots indicate the number of siRNA target genes identified in the immunoprecipitated sRNA dataset by the authors’ analysis (WAGO-4 IP, CSR-1 IP) and/or by smalldisco.

We validated the siRNA counting function using publicly available datasets from Scheer *et al*. (2021). This study sequenced sRNAs from wildtype and three mutant strains of *Arabidopsis* and identified siRNAs within those four datasets. The authors provide a brief description of how they identified siRNA reads wherein their method uses logic similar to ours except that siRNAs were not required to map antisense to coding regions. We evaluated the four datasets with smalldisco and compared the output to the authors’ results. For each genotype, the number of siRNA reads identified by smalldisco was similar to that identified by the authors (Figure 3). Compared to the authors’ analysis, smalldisco identified: 5% more siRNA reads in wildtype, the same number of reads in the *urt1-1* single mutant, 88% as many reads in the *xrn4-3* single mutant, and 85% as many reads in the *urt1-1 xrn4-3* double mutant. We note that our approach of identifying antisense reads uncovered a similar number of siRNAs as did the authors’ approach of requiring matching sense and antisense reads.

**Figure 3.**
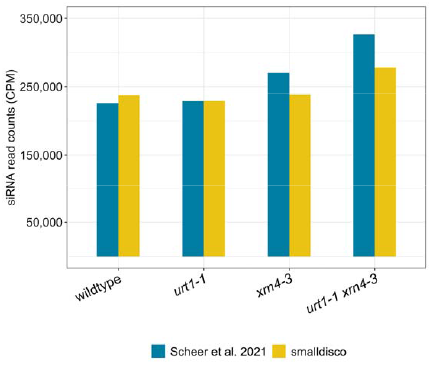
Identification of siRNA reads in small RNA sequencing datasets. SiRNA read counts are plotted as counts per million reads for small RNA sequencing datasets obtained for wildtype and three mutant genotypes. Blue bars indicate read counts reported by Scheer *et al*. (2021) and yellow bars indicate reads identified by smalldisco.

One published protocol for identifying siRNAs and for which software is available is segmentSeq (Hardcastle *et al*. 2011). segmentSeq assumes all siRNAs are generated by Dicer endonuclease cleavage of double-stranded RNA, and therefore reads matching the sense strand should be present in the dataset. Although most *C. elegans* siRNAs are generated as direct single-strand RNA products of RNA-directed RNA polymerase and do not require Dicer activity, we were interested in comparing sRNA discovery by the two programs. We analyzed the Davis *et al*. (2018) dataset (see Methods) with each program and noted several major differences in output. First, smalldisco analyzed our sample input dataset in ~2.5 hr, whereas segmentSeq required ~2.5 weeks. Second, the two programs identified a common set of 6830 sRNA-producing genes and another smaller but substantial number of sRNA producing genes that are unique to one or the other program (1835 for segmentSeq, 3899 for smalldisco) (Figure 4A). Third, the two programs mapped sRNAs to the genome—and hence identified siRNA-associated genes—at different spatial scales. smalldisco identified an average siRNA region of 124.3 bp (± 7.4 bp SE) when run with the default “CDS” (exon) setting, whereas segmentSeq identified an average siRNA region of 6.7kb (± 68.9 bp SE) corresponding, in some cases, to more than one gene (Figure 4B, Figure S1). Although segmentSeq is a useful tool for siRNA identification, smalldisco gives finer grained results and is substantially faster.

**Figure 4.**
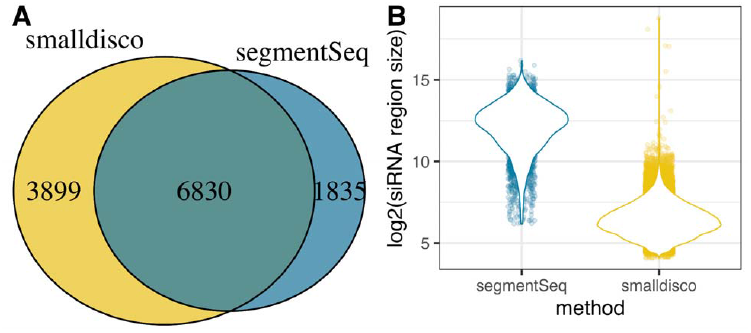
siRNA prediction comparison between smalldisco and segmentSeq. (A) Overlap between the number of sRNA target genes predicted using smalldisco and segmentSeq. Note that segmentSeq is not specific to siRNAs, but presumably most sRNAs derived from coding sequences are likely to be siRNAs. (B) The two methods differ substantially in the size of their predicted siRNA region, with smalldisco having an average size of 124.3 bp (±7.4 bp SE) and segmentSeq having an average size of 6.7kb (±68.9 bp SE).

In summary, smalldisco is a command-line tool for siRNA discovery and annotation from small RNA sequencing datasets. We expect that smalldisco will be useful in identifying, annotating, and quantifying the abundance of siRNAs in wildtype and mutant datasets derived from whole animals and dissected tissue. Inclusion of the Tailor program in the pipeline allows the user to identify and quantify 3’ non-templated nucleotides that may be present on the siRNAs or any other small RNA in the dataset.

## Data availability

smalldisco and supporting documentation is available on GitHub (https://github.com/ianvcaldas/smalldisco).

## Acknowledgements

The authors thank Sarah Hall and two reviewers for helpful comments on the manuscript.

---

**Figure S1.**
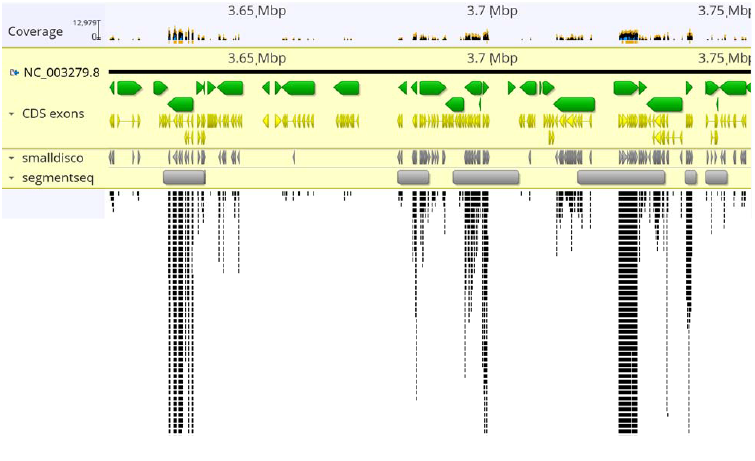
Examples of siRNAs identified by smalldisco and segmentSeq that map to one region of the genome. Image is a screen shot from the Geneious genome browser. Green bars indicate the positions of transcribed genes, with the pointed end indicating direction of transcription. Yellow bars indicate coding exons. Gray bars indicate siRNAs as identified by smalldisco (top) and segmentSeq (bottom). Stacked black lines represent siRNA-derived read pileups.

**Table S1:**
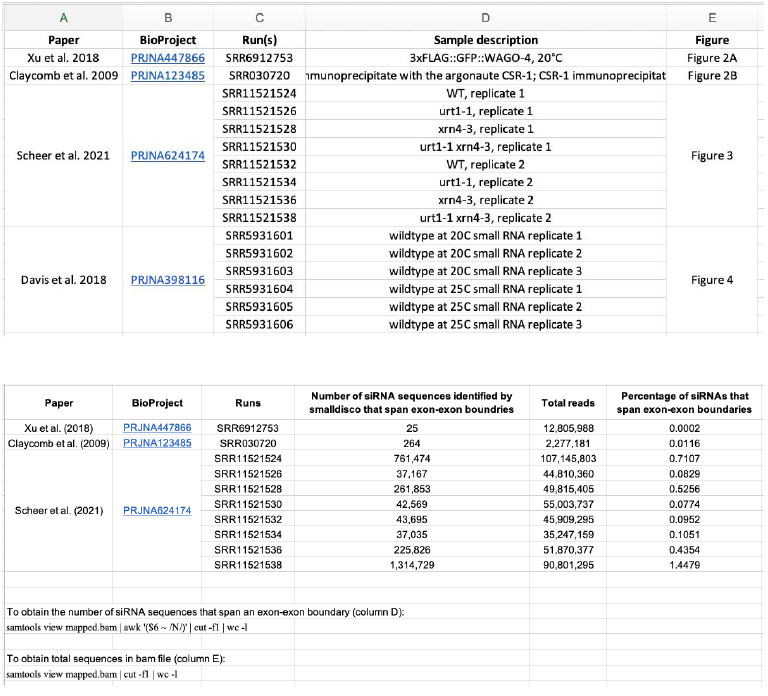
List of NCBI accession numbers used in this study and frequency of siRNAs that map to exon-exon boundaries.

