## Supplementary figures and images for "smalldisco, a pipeline for siRNA discovery and 3’ tail identification"

### Figure S1

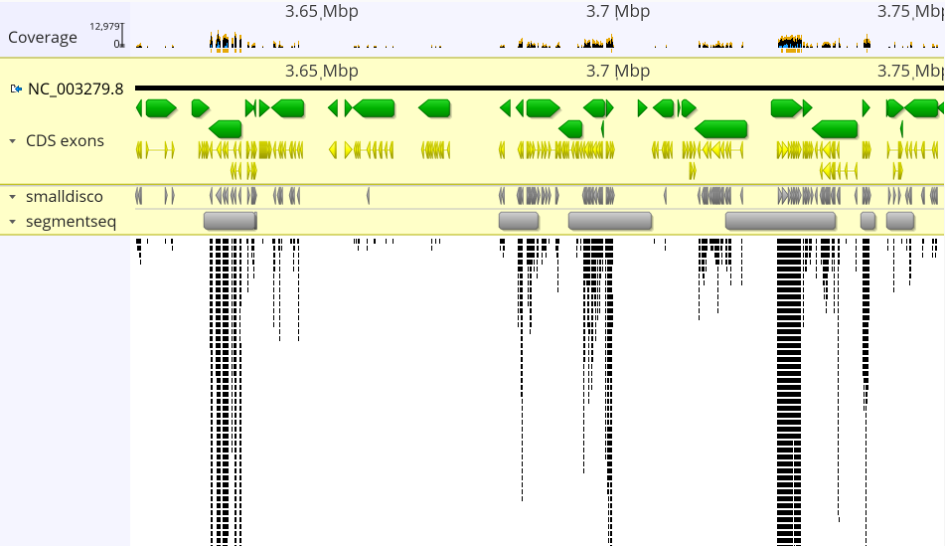
